# ColorSTEM: An easy and convenient approach to Multicolor Electron Microscopy of Labeled Biological Specimens

**DOI:** 10.64898/2025.12.15.694448

**Authors:** Ranjan Ramachandra, Mason R. Mackey, Junru Hu, Tristan M. Shone, Mia V. Flores, Isabella A. Ramos, Steven T. Peltier, Stephen R. Adams, Mark H. Ellisman

## Abstract

A new method for Multicolor Electron Microscopy (EM) using scanning transmission EM, ColorSTEM has been developed to increase the sensitivity, instrument availability and ease of use compared to that previously developed using electron energy loss spectroscopy and energy filtered transmission EM. Both methods paint multiple subcellular complexes with different colors, enabled by the localized and targeted deposition of diaminobenzidine (DAB) conjugated to different metals, which provides the differentiating signal from each other. With ColorSTEM, elastic scattering that is dependent upon atomic weight (Z-contrast) has far greater signals. Using deposited DAB with copper or gold to label subcellular components, Z-contrast imaging generates a pseudo-elemental map, resulting from differences in the grayscale of the conventional STEM images.

## INTRODUCTION

Multicolor fluorescence microscopy is a mainstay of current cell biology and biomedical research as it enables rapid monitoring of specific cellular components (proteins, lipids, organelles etc.) in living and fixed biological samples ^1, 2^. However, it is limited in the number of entities that can co-measured by the spectral width of the fluorophores, with the vast bulk of cellular components and compartments being invisible. In comparison, electron microscopy (EM) of biological samples provides far superior resolution and detail of the cellular ultrastructure, unmatched by fluorescence microscopy despite the many recent advances in its techniques ^3^. The standard technique of electron microscopy of biological samples involves staining the cellular components including proteins, lipids, RNA, and DNA with heavy metals such as osmium, lead and/or uranium, which enable acquisition of high-resolution greyscale EM images. Although electron microscopy can provide complex 3-dimensional structural details of the cellular subcomponents with much clarity, it lacks the context provided by different fluorescent labels as in Multicolor Fluorescence microscopy.

A recent advance, MultiColor EM, combines labeling and selective staining of multiple subcellular components with advanced EM imaging to overcome this limitation, enabling fluorescence-like color images while retaining the full resolution of EM ^4^. The Multicolor EM technique involves sequential deposition of diaminobenzidine (DAB) conjugated to different lanthanide chelates (Ln-DAB2) at specified proteins or cellular targets by photosensitizers, small-molecule probes, or peroxidases. The lanthanides typically used are lanthanum (La), cerium (Ce), praseodymium (Pr) and neodymium (Nd). Their distinct elemental maps are obtained by energy-filtered transmission electron microscopy (EFTEM) using an electron energy loss spectrometer (EELS), which immediately validates the localization and structure of the targeted protein or macromolecule. The color-coded elemental map(s) overlaid over the conventional EM image, generate the Multicolor EM image. This technique has been used to answer many biological structure-function questions, which otherwise would not have been possible ^4–8^.

At the characteristic edge or core-loss for the lanthanide metals, the elemental signal is an extremely small fraction of the incident beam, and the acquisition of their elemental maps is often challenging. As a first step to improve the poor signal-to-noise (SNR) in the elemental maps, we demonstrated the use of Direct Detection Device (DDD), instead of the traditional phosphor based scintillator imaged by a photosensitive Charged Coupled Device (CCD), which provided a 3X boost in SNR ^9^. Alternatively, it was proposed that acquiring elemental maps at the intermediate-loss energy (99 – 120 eV), where the signal is substantially higher than the core-loss energy (800 – 1000 eV) should circumvent the poor SNR in elemental maps ^10^. However, over these intermediate-loss energies, although the SNR is good, the signal above background can be very low. What’s more, since these edges lack a sharp and clear shape (vs. core-loss edges), they can often lead to artifacts in the computed elemental maps ^10^.

MultiColor EM of biological samples has also been achieved by using Energy Dispersive X-ray spectroscopy (EDX) in a scanning electron microscope ^11^; however, EDX is generally considered a lower resolution technique in comparison to EELS ^12^. Recently, there have been tremendous progress in the instrumentation of EDX detectors, and single atom elemental maps have been acquired ^13, 14^. These maps have mostly been acquired of semiconductor/material science samples, which are usually less prone to sample drift and beam induced specimen damage compared to biological samples.

Despite the improvements made in both the EELS and EDX techniques, acquisition of elemental maps is somewhat involved, and both methods require expensive instrumentation and highly skilled personnel to operate them. In this paper, we describe a method to create the MultiColor EM images without the need for either the EDX or EELS instrumentation, by utilizing the scanning transmission electron microscopy (STEM), which generally comes as a standard package with most 200 keV and 300 keV TEM.

The ‘atomic-number contrast’ or ‘Z-contrast’ is the prominent mode of imaging in STEM ^15–22^ and is due to the Coulombic attraction of the incident electron beam and partially screened nucleus of the target atom and thus proportional to atomic number Z of the sample ^23, 24^. In fact, one of the earliest elemental maps in an electron microscope was acquired using Z-contrast imaging to discern the grey scale difference between platinum and palladium atoms ^21^. It should be emphasized that a Z-contrast image is not a true elemental map like an EELS/EFTEM or EDX map, but rather, if there is *a priori* information of the metals present in the sample, then subtle differences in their greyscale pixel values can be used to create pseudo-color elemental maps. Therefore, although the STEM Z–contrast maps are not as robust as EFTEM/EELS elemental maps, the trade-off is the ease and convenience of use. Since STEM is more readily available in most EM laboratories, this will enable more widespread adoption of the Multicolor EM labeling technique by the scientific community.

## MATERIALS AND METHODS

### Samples

#### Single labeling of cells with gold or copper

##### Cell preparation and fixation

HEK293T cells were plated onto MatTek dishes containing 35mm glass bottom No. 0 coverslips coated with poly-d-lysine (P35GC-0-14C, MatTek Corporation). Three days later cells were incubated with 10 μM EdU (CCT-1149, Vector Laboratories for 18 hours. The cells were fixed with 2% EM grade glutaraldehyde (18426, Ted Pella Incorporated) in PBS, pH 7.2 for 5 minutes at 37oC and then on ice for 55 minutes. Cells were washed (5 x 1 min) with PBS on ice, blocked with 20 mM glycine in PBS for 20 minutes on ice and washed (2 x 1 min) with PBS buffer on ice.

##### Click reaction of Fe-TAML-azide to EdU-labeled cells

The click reaction of the EdU-fed cells was carried out at room temperature and protected from light by incubation with a freshly prepared mixture of 28 μM Fe-TAML azide ^8^ in 900 μl click buffer (100 mM NaCl 50 mM Na·HEPES pH 7.6, 0.1% Saponin (Sigma, S-4521)) and 10.0 μl CuSO4 (100 mM in water). The reaction was initiated by adding 100 *μ*l of freshly prepared aqueous sodium ascorbate (100 mM). After 30 minutes, a second 100 *μ*L aliquot of newly prepared aqueous sodium ascorbate (100 mM) was added for a further 30 minutes incubation. Cells were washed with PBS pH 7.4 (5 x 1 min) at room temperature and washed with 100 mM NaCl 50 mM Na·Bicine pH 8.3 (5 x 1 min).

##### Cu-DAB2 deposition with Fe-TAML catalyzed oxidation by H_2_O_2_

2 mM Cu-DAB2 solutions in 100 mM 100 mM NaCl 50 mM bicine pH 8.3 were prepared similarly to previously reported solutions of Ln-DAB2 in cacodylate buffer ^4^ by substituting with stock solutions of 0.1 M CuSO4, 1 M Na·bicine pH 8.3 and 1 M-NaCl. 10 μl of H_2_O_2_ from a 30% stock solution was added to 10 ml of the 2 mM Cu-DAB2 solution. The Cu-DAB2 reaction ran for one hour on a set of cells and then washed with bicine buffer (5 x 1 min) on ice and processed for TEM.

##### DAB deposition by Fe-TAML catalyzed oxidation by H_2_O_2_

5.4 mg of 3,3’-Diaminobenzidine (DAB) (D8001-10G, Sigma-Aldrich) was dissolved in 1.0 ml of 0.1 N HCl and 9.0 ml of 50 mM Na·bicine 100 mM NaCl pH 8.3 was added with 10 μl H_2_O_2_ (final, 40 mM from 30% stock) to the DAB solution. The DAB/H_2_O_2_ solution was added to a set of cells at room temperature and the reaction time was 20 minutes. Cells were washed with 100 mM NaCl 50 mM Na·Bicine pH 8.3 (5 x 1 min).

##### Gold intensification of precipitated DAB

Gold intensification of DAB followed previously reported methods ^25–27^. Cells were washed (3 x 1 min) with 2% sodium acetate, pH 7.4 on ice and then prewarmed at 60°C for 10 min. The enhancement solution containing 3% hexamethylenetetramine in ddH2O, 5% silver nitrate in ddH2O, and 2.5% disodium tetraborate in ddH2O, mixed in a ratio of 20:1:2, prewarm solution for 10 minutes at 60oC, was added to the cells and incubated for 8 minutes at 60°C. The cells were washed (5 x 3 min) with 2% sodium acetate pH 7.4 at room temperature and then incubated with 0.05% gold (III) chloride trihydrate (Sigma-Aldrich, 520918) in ddH2O for 5 minutes at room temperature. After washing (5 x 3 min) with 2% sodium acetate pH 7.4 at room temperature, the cells were incubated with 2.5% sodium thiosulphate in ddH2O for 4 min at room temperature, washed (1 x 3 min) with 2% sodium acetate pH 7.4 at room temperature, with PBS pH 7.2 (5 x 1min) and processed for TEM.

##### Double labeling of cells with gold and copper

HEK293T cells were cultured on imaging plates containing poly-d-lysine coated glass bottom No. 0 coverslips (P35GC-0-14C, MatTek Corporation). Three days later cells were incubated with 10 μM EdU for 12 hours. Cells were also transiently transfected with either miniSOG-mitomatrix fusion or miniSOG-KDEL fusion using Lipofectamine 3000 (Life Technologies). The next day the cells were fixed with 2% EM grade glutaraldehyde (18426, Ted Pella Incorporated) in PBS pH 7.2 (18851, Ted Pella Incorporated) for 5 minutes at 37° C and then on ice for 55 minutes. Cells were washed (5 x 1min) with PBS on ice, blocked with 20 mM glycine in PBS for 20 minutes on ice, and washed (2 x 1 min) with PBS buffer on ice. For miniSOG transfected cells, regions of interest were identified by standard GFP filter set (ex475) at light intensity of 10% and for photooxidation of DAB in 1X PBS at pH 7.2 the light intensity was set at 100% using a Leica SPEII confocal microscope. The photooxidation reaction occurred for 2-3 minutes to give the desired brown intensity color from the precipitate. After photooxidation, gold intensification of DAB method was followed. The click reaction for EdU with Fe-TAML azide was then carried out as described above. The cells were then washed (5 x1 min) with 100 mM NaCl 50 mM Na·bicine pH 8.3. H_2_O_2_ (10 µL of 30% stock solution) was added to 10 mL of 2.5 mM DTPA-DAB2 ^28^ in 100 mM NaCl 50 mM Na·bicine 2.5% DMF pH 8.3 and the solution was added to the cells for 30 minutes to produce the desired brown intensity color from the precipitate. Cells were then washed (5 x 2 min) with 25 mM acetic acid pH 5.6 and 50 mM CuSO4 in 25 mM acetic acid pH 5.6 was added to the cells for 30 min. Cells were then washed (5 x 2 min) with 25 mM acetic acid pH 5.6 on ice.

##### EELS acquisition on astrocyte filled with CeDAB2

Hippocampal astrocytes from a perfused mouse (2-month-old BALB/c male) were intracellular filled with Lucifer Yellow and Neurobiotin/Alexa 568 ^29^, treated with blocking buffer for 15 minutes. The Lucifer Yellow was photooxidized and Neurobiotin/Alexa 568 was enzymatically polymerized, a detailed description is already published ^4^.

##### TEM processing

Cells were washed with (5 x 1 min) with cold dd H_2_O on ice, and dehydrated in 20, 50, 70, 90 and 100% aqueous ethanol on ice for 1 minute each. Further dehydration (3 x 2 min) with 100% ethanol at room temperature was followed by infiltration with a 1:1 ratio of 100% Durcupan ACM resin (14040, EMS) and 100% dry ethanol for 30 minutes, change to 100% Durcupan epoxy resin overnight. The 100% Durcupan resin was changed (3 x 1 hour) and placed in a vacuum oven 60°C to be embedded.

##### Sectioning

MatTek dishes were taken out of the vacuum oven, the labeled cells were identified, cut out of the dishes and glued on to plastic dummy blocks. Coverslips were removed and all specimen blocks were sectioned at 100 nm using a Lieca Ultracut ultramicrotome with an Ultra 450 Diatome diamond knife and sections were picked up on copper 50 mesh grids (G50, Ted Pella Incorporated). The sections on the grids were carbon coated on both sides of the section using a Cressington 208 carbon coater.

### Electron Microscopy

The EFTEM/EELS work was performed using a Thermo Scientific Talos 200 keV equipped with a CFEG source. The microscope was fitted with a Gatan Continuum GIF Spectrometer for EELS and EFTEM. A 2k X 2k CMOS based Continuum camera was used to record the images and spectra. All images and spectra were recorded with a condenser aperture of 150 µm, objective aperture of 100 µm, entrance aperture of 5 µm and collection semi-angle of 100 mrad. Spectra were collected at an energy dispersion of either 0.3 eV/pixel or 0.75 eV/pixel. The spectrometer energy resolution was 1.5 eV, measured as the FWHM of the zero-loss peak obtained at a Spectrum dispersion of 0.3 eV. Before the acquisition of EELS spectra and STEM datasets the sample was pre-irradiated with a low beam dose of ∼ 3.5 × 10^4^ e/nm^2^ for about 30 mins to stabilize the sample and to reduce contamination ^30^.

The STEM datasets were acquired using a FEI Titan at 300 keV, equipped with a XFEG source. All the images were acquired on a Fischione High Angle Annular Dark Field (HaaDF) detector. The collection angle of the detector was varied by changing the camera length – the smaller the camera length, the larger the collection angle (See Table 1). All the datasets were collected with a probe-forming condenser aperture of 50 µm, probe semi-angle of 6.7 mrad, spot size of 5 and probe current of 55 pA. Samples imaged in STEM mode exhibit extreme tolerance to the total electron dose they can withstand without significant damage, generally several orders of magnitude higher than in TEM mode ^30, 31^; however, they are extremely sensitive to the dose rate and, beyond a threshold, the samples crack easily. To avoid beam damage, the pixel dwell time was set to 6 µsec (2.2 e/nm^2^.sec), but the SNR of a single STEM image at this dose rate was low, specifically for the larger collection angle images which are signal limited. To increase the SNR, multiple images were acquired at each camera length, aligned using the Template matching plugin in ImageJ ^32^, and then summed. The total signal in an image increased, going from smaller (44 mm) to larger camera lengths (430 mm), allowing for less image frames to be acquired at the larger camera lengths. The summed images were then divided by total number of image frames for that particular collection, thereby normalizing the signal to a single image but with a much higher SNR. At each of these settings, 10 dark images (i.e. images with column valves closed) were collected, and the average was subtracted from the normalized summed image. All STEM datasets (i.e., multiple STEM images at each of the camera lengths) were acquired by an automatic subroutine written in Serial EM ^33^. The subroutine mainly alters the brightness and contrast of the STEM detector for the different camera lengths. The routine also automatically adjusts the diffraction alignment to bring the beam to the center of the HaaDF detector. The brightness and contrast of the STEM detector was adjusted for each camera length to get the maximum dynamic range without oversaturating the pixels. Also, between each successive STEM image acquisition, a low dose of sample irradiation i.e., beam showering was performed, to mitigate effects of sample charging and hydrocarbon contamination ^30, 34^.

**Table 1:** The collection angles for the different camera length for the FEI Titan microscope operated at 300 keV.

| Camera Length<br>in mm | Inner collection<br>Angle ($\Theta_1$) mrad | Outer collection<br>Angle ($\Theta_1$) mrad |
| --- | --- | --- |
| 44 | 132 | 200 |
| 55 | 106 | 200 |
| 69 | 84.1 | 200 |
| 87 | 66.7 | 200 |
| 110 | 52.8 | 200 |
| 135 | 43 | 200 |
| 175 | 33.2 | 200 |
| 220 | 26.4 | 161 |
| 275 | 21.1 | 129 |
| 340 | 17.1 | 104 |

The Z-contrast or elemental STEM map was generated by computing the rate of change in the signal intensities for the different elements at the different camera lengths. In this paper, the images acquired at camera lengths of 44 mm, 55 mm and 69 mm were found sufficiently adequate to extract the elemental signal. The rate of change of signal intensities between camera lengths varied from one pixel to another by virtue of the elemental composition present in the pixel, and this variation was used to extract the elemental signal using the simple single line command “tert” available in Digital Micrograph.

All the SEM imaging was done on a Zeiss Merlin SEM operated at high vacuum. The SEM was generally operated at energies of 3 - 7 KeV beam energy, with the simultaneous utilization of the in-built Zeiss deceleration stage, capable of providing uniform and consistent negative bias on the stage from 0 – 5 KeV. All images were acquired on the Zeiss Sense BSD detector. The Samples were thin sections prepared for TEM/STEM, mounted on a STEM-in-SEM kind of holder.

## RESULTS AND DISCUSSION

To demonstrate the poor sensitivity of a MultiColor EM image, we used a hippocampal section with individual astrocytes injected with Lucifer Yellow. Ce-DAB2 was polymerized by photooxidation, and Ce detected at the M_4,5_ edge occurring at 883 eV ^4^. Figure 1A, a TEM of a Ce-labelled astrocyte is shown with its EELS spectrum taken from 0 eV to 2800 eV (Figure 1B). The sharp needle like peak appearing at 0 eV, is the zero-loss peak, which mostly contains the elastically scattered electrons that have suffered no energy loss or very negligible energy loss. The small bump next to the zero-loss peak is the plasmon peak, which are comprised of electrons that have undergone minimal energy loss due to interactions with the weekly bound outer shell electrons of the sample. The signal drops exponentially after the plasmons, generally the strongest elemental edge in biological samples is observed for carbon at ∼ 284 eV, but this edge is not even visible in the spectrum shown. The region of the spectra enclosed in the rectangular box is magnified (Figure 1B’) to show the carbon edge at 284 eV but the cerium edge at ∼ 883 eV is still not discernible. This could only be captured by operating the EELS spectrometer at a much higher energy dispersion (i.e. energy resolution measured in eV/pixels) and acquiring the spectrum for much longer exposure time to reveal the double Ce M_4,5_ edge (Figure 2C). The signal for each the peak/edge i.e. zero-loss peak (0 eV– 20 eV), Plasmon (20 eV – 50 eV), carbon edge (285 eV – 315 eV) and cerium edge (885 – 915 eV) can be integrated and compared to the total incident electron beam. The zero-loss peak and the plasmon contain 74.3% and 17.6% of the total incident beam intensity respectively, together they contain 91.9% of the total beam intensity. In comparison, the carbon edge and the cerium edge contain 0.1% and 0.014% of the total beam intensity respectively, i.e. there is ∼ 1 electron in the cerium edge detected for every 10000 electrons incident on the sample.

**Figure 1.**
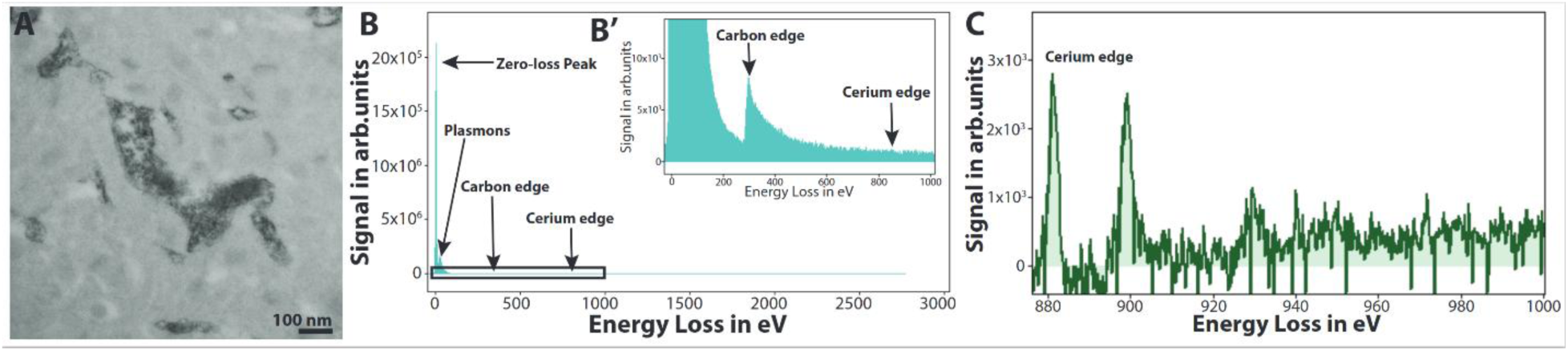
**A**, Astrocyte in a mouse hippocampal slice injected with Lucifer Yellow and Ce-DAB2 polymerized by photooxidation. **B**, EELS spectra from 0 eV - 2800 eV acquired at 1.5 eV per pixel for an exposure time of 2 x 10^-5^ sec per frame and summed for 2000 frames. **B′**, expansion of boxed rectangular region of the spectra. **C**, EELS spectrum of A acquired at an spectrum dispersion of 0.15 eV per pixel for an exposure time of 24 sec, summed for 6 frames, showing the presence of cerium.

**Figure 2.**
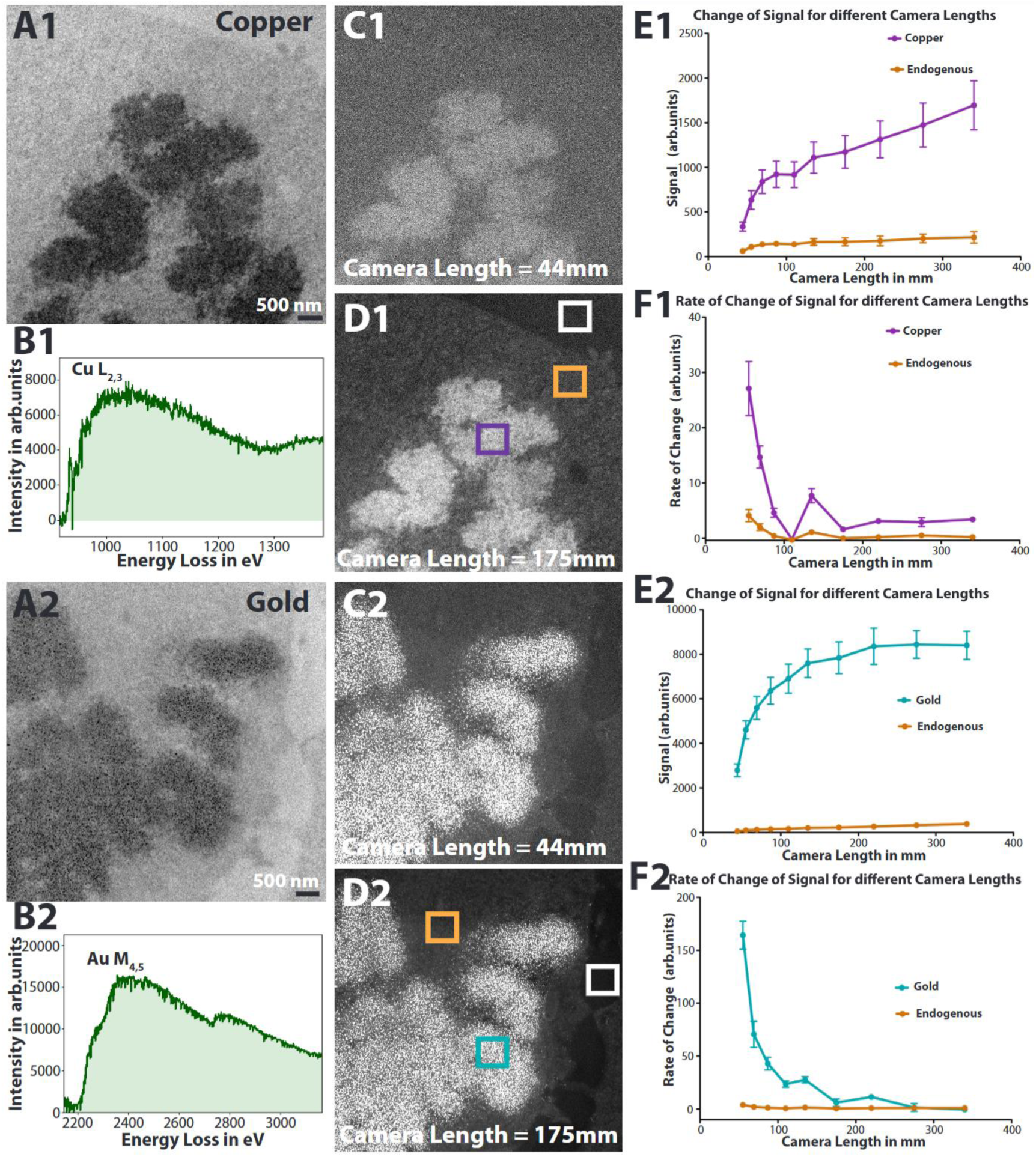
STEM datasets acquired on HaaDF detector at different camera lengths (i.e. different collection angles) of mitotic HEK cells with condensed chromosomes labeled with precipitated DAB bearing Cu or Au. DNA was click labeled with FeTAML-azide to metabolically incorporated EdU. **A1**, Conventional TEM image showing the mitotic cell with the Cu metal stain on the DNA. **B1**, EELS spectrum confirming the presence of Cu in the sample. **C1**, STEM HaaDF image collected at the smallest camera length of 44 mm i.e. the largest scattering semi-angle of 132 to 200 mrad. The image looks noisy due to limited signal, since fewer electrons are scattered to these large angles. **D1**, STEM HaaDF image collected at an intermediate camera length (175 mm) i.e. the scattering semi-angle of 33.2 to 200 mrad. The white colored box in the image, shows a region in the sample that contains only resin, all the HaaDF STEM images were normalized to this signal at the respective camera length, by its subtraction. The blue and the orange box in the image is where the signal from Cu and endogenous cellular material were measured respectively. **E1**, Change in signal intensities for Cu and the endogenous cellular material as a function of camera length. **F1**, rate of change in signal intensities, i.e. slope of the graph shown in **E1**, for Cu and the endogenous cellular material as a function of camera length. **A2**, Conventional TEM image showing the mitotic cell DNA labeled with Au metal. **B2**, EELS spectrum confirming the presence of Au in the sample. **C2**, STEM HaaDF image collected at the smallest camera length (44 mm) i.e. the largest scattering semi-angle of 132 to 200 mrad. The image does not look as noisy as in C1, because scattering cross-section is proportional to Z^2^, and Au scatters more electrons into the larger scattering angles than Cu. **D2**, STEM HaaDF image collected at an intermediate camera length (175 mm) i.e. the scattering semi-angle of 33.2 to 200 mrad. The white colored box in the image, shows a region in the sample that contains only resin, all the HaaDF STEM images were normalized to this signal at the respective camera length, by its subtraction. Blue and orange boxes are areas where the signal from the Au and the endogenous cellular material was measured respectively. **E2**, change in signal intensities for Au and the endogenous cellular material as a function of camera length. **F2**, rate of change in signal intensities, i.e. slope of the graph shown in E1, for Au and endogenous cellular material as a function of camera length.

As evident from this example, the EFTEM technique of acquiring elemental maps for Multicolor EM is inherently a low SNR problem, requiring acquisitions with very high total dose and dose rates. In comparison, the ‘Z-contrast’ imaging in STEM, which is due to the elastically scattered electrons can be as high as 10-20 % of the total incident beam intensity ^35–37^. Z-contrast imaging has been able to resolve individual boron and nitrogen atoms in a single atomic monolayer of boron nitride, using an aberration corrected STEM ^38^. The fact that boron (Z = 5) and nitrogen (Z = 7), being just two atomic numbers separated could be clearly distinguished from each other proves the validity of the technique. However, for stained biological plastic sections, homogenous samples of atomic thickness are not practical. Also, for Multicolor EM samples, in addition to the lanthanide metals, there are endogenous elements like C, N, O, P, S and the elements from exogenous stains/fixatives like As, Os, Pb, U, all of which may be attached or combined with each other, further confounding the Z-contrast signal from the lanthanide metals. Therefore, distinguishing the signal from the two lanthanide metals in a Multicolor EM sample that are only 1 or 2 atomic numbers apart by Z-contrast STEM imaging is rather improbable. However, if the two metals bound to the DAB are well separated in the periodic table (i.e., many atomic numbers apart), and influence from other exogenous stains/fixatives can be minimized or eliminated, then the Z-contrast signal differential from these two metals may be large enough that their discrimination is plausible.

To test this hypothesis, Multicolor EM tags were developed for two metals with a large Z separation, Cu (Z = 29) and Au (Z = 79) respectively. Cu was bound to DTPA-DAB2 to form Cu-DAB2 and DAB was intensified with Au using known procedures. Z-contrast signal was first measured on samples containing only a single metal instead of two metals to avoid any contamination due to the unwanted deposition of the second metal at the sites where the first metal is bound ^4^. Samples were prepared by click-labeling of DNA that had metabolically incorporated EdU with the small molecule peroxidase, FeTAML-azide, followed by deposition of Cu-DAB2 or DAB enhanced by Au. The samples were prepared with rigorous detail, to avoid adding any heavy elements in the sample by the way of reagents/ buffers used (such as arsenic), and without post-staining with osmium or other heavy metals.

Figure 2A1 and Figure 2A2 are conventional TEM images of Cu and Au labeled DNA, respectively in HEK cells. To calibrate their Z-contrast signal intensities, mitotic cells undergoing cell division was specifically chosen to concentrate the signal and increase its homogeneity, thereby providing better precision in the measurement of the signal. Figure 2B1 and Figure 2B2, are the EELS spectra confirming the presence of Cu and Au respectively with edges at 931 and 2206 eV respectively. Z-contrast images were acquired on a HaaDF detector, which is an annular ring detector with a hole at the center, and the collection angle (i.e. the angular range for which electrons elastically scattered from the sample are accepted by the detector) varied by changing the camera length of the detector. The collection angles for the different camera length for the FEI Titan microscope operated at 300 keV are shown in Table 1.

To achieve Z-contrast imaging in STEM, one of the primary requirements is that the imaging conditions should predominately be incoherent ^16, 21^. If the HaaDF detector collects coherent Bragg diffracted beams along with the Z-contrast signal, image interpretation becomes rather confusing ^19^. Increasing the inner collection angle (Ɵ_1_) of the HaaDF detector tends to collect electron beams with larger scattering angles, which should reduce the collection of diffracted beams. Also, channeling contrast, which occurs when the electron beam is preferentially scattered along or between crystal planes, exists at much higher scattering angles than Bragg peaks, and Ɵ_1_ > 100 mrad is recommended to avoid these deleterious effects to Z-contrast imaging ^17, 20, 39, 40^. For plastic embedded biological samples, which are mostly amorphous in nature, it has been suggested that incoherent imaging conditions should prevail at Ɵ_1_∼>15 mrad or so ^35^. Generally, it is accepted that setting the inner collection angle Ɵ_1_ > 40–50 mrad, should facilitate incoherent imaging conditions, with the rule of thumb being Ɵ_1 >_ 3Ɵ_0_, Ɵ_0_ being the probe forming angle (8.7 mrad for this experiment) ^16, 22^.

Figure 2C1 and Figure 2C2 are the STEM HaaDF images collected at a camera length of 44 mm, corresponding to the largest collection semi-angle of 132 to 200 mrad, of Cu and Au labeled DNA respectively. Figure 2C2 does not look as noisy as in Figure 2C1, because the scattering cross-section is proportional to Z^2^, and gold scatters far more electrons into the larger scattering angles than Cu. This is the basis for Z-Contrast imaging in STEM. Figure 2D1 and Figure 2D2 are the corresponding STEM HaaDF images collected at a camera length of 175 mm, an intermediate collection semi-angle of 33.2 to 200 mrad, of Cu and Au respectively, where the incoherent imaging condition of Ɵ_1 >_ 3Ɵ_0_ is still valid. At each camera length multiple image frames were collected and processed as explained in detail in the methods section. The white colored box in Figures 2D1 and 2D2 shows a region that contains only the resin, the HaaDF STEM images were normalized to this signal at the respective camera length, by its subtraction. The blue box where the signal from the endogenous cellular material was measured, and the orange box is where the signal from the Cu or Au signals were measured. Figure 2E1 shows the change in signal intensities for Cu and the endogenous cellular material as a function of camera length and in Figure 2E2, the corresponding change for Au and the endogenous cellular material. Figures 2E1 and 2E2 demonstrate how electron beam scattering by Cu and Au atoms influences the signal captured by the HaaDF detector at different collection angle settings. Although the trend of signal intensity change with camera length (i.e. collection angle) is important, the actual values obtained can be misleading as they are not just dependent on the scattering angles but also are a function of the brightness and contrast values used to capture to the image. A more unbiased measurement is the rate of change of signal (i.e, the slope of the change in signal with respect to camera length) with change in collection angle. Figure 2F1 the rate of change of signal for Cu and the endogenous cellular material is plotted as a function of camera length; Figure 2F2 shows the corresponding rate of change of signal for Au and the endogenous cellular material. From Figures 2F1 and 2F2, it can be seen that the rate of change of signal for endogenous cellular material varies from ∼5 to 0 and is consistent for both the cases whereas the rate of change of signal for Cu varies from ∼27 to 0 and that for Au it varies from ∼153 to 0. The differential between the rate of change of signal between Cu and Au is the most at the largest collection angles, consistent with Z-contrast being more prevalent at the larger collection angles. When the camera length was changed from 44 mm (Ɵ_1_ = 132 mrad) to 55 mm (Ɵ_1_ = 106 mrad), the rate of change in Cu and Au signal was ∼27 and 153 respectively. Changing the camera length from 55 mm (Ɵ_1_ = 106 mrad) to 69 mm (Ɵ_1_ = 84.1 mrad), the rate of change in Cu and Au signal is ∼15 and 46 respectively. There is therefore a substantial difference between the rate of change in signal between Cu and Ar for camera lengths 44 mm (Ɵ_1_ = 132 mrad) to 135 mm (Ɵ_1_ = 43 mrad). At a camera length of 175 mm (Ɵ_1_ = 33.2 mrad) and higher this differential is minimized, which also coincides with the limit for incoherent imaging (i.e. Ɵ_1 >_ 3Ɵ_0_). Therefore, it is the signal and its rate of change at camera lengths 44 mm, 55 mm and 69 mm, corresponding to the largest collection angles that are critical to extracting the elemental maps.

Figure 3 demonstrates the application of Z-Contrast enabled ColorSTEM imaging on the Multicolor EM samples prepared by labeling of DNA with Cu in HEK cells by click reaction of FeTAML-azide to EdU as above. Endoplasmic reticulum (ER) was labeled with Au by transient transfection of miniSOG-KDEL, light-driven DAB precipitation and its enhancement with Au. With no post-staining the only metals in the sample were Cu and Au. Figure 3A is a conventional TEM image showing the Cu metal stain on the DNA and the Au metal stain on the ER. Figures 3B and 3C show the EELS spectra confirming the presence of Cu and Au in the sample, respectively. Figures 3D and 3E show the STEM HaaDF image collected at the camera length of 44 mm (Ɵ_1_ = 132 mrad) and 175 mm (Ɵ_1_ = 33.2 mrad), respectively. The white box in Figure 3D, is a region in the sample that contains only resin; all the HaaDF STEM images at each camera length were normalized to this signal. The cyan, magenta, and orange boxes in Figure 3D are where the signals from Au, Cu and the endogenous cellular material were measured respectively. Figures 3F and 3G show the average change in signal intensities and rate of change in signal for the Au, Cu and the endogenous cellular material in five different regions as a function of camera length, respectively. The trend in rate of change of signal for Au, Cu and endogenous cellular material for both the single color (Figures 2E2 and 2F2) and the multicolor samples (Figure 3G) are very similar and corroborate each other. From Figure 3G, the rate of change of signal for Au from camera length 44 mm (Ɵ_1_ = 132 mrad) to camera length 55 mm (Ɵ_1_ = 106 mrad) is 303.1 ± 50.8; and from camera length 55 mm (Ɵ_1_ = 106 mrad) to camera length 69 mm (Ɵ_1_ = 84.1 mrad) is 92.3 ± 21.7. The Au elemental map (Figure 3H) was computed by picking pixels whose rate of change of signal going from camera lengths 44 mm to 55 mm was >100, and going from camera lengths 55 mm to 69 mm was >35. The rate of change of signal for Cu from camera length 44 mm (Ɵ_1_ = 132 mrad) to camera length 55 mm (Ɵ_1_ = 106 mrad) is 50.9 ± 6.5; and from camera length 55 mm (Ɵ_1_ = 106 mrad) to camera length 69 mm (Ɵ_1_ = 84.1 mrad) is 25.4 ± 3.2. The Cu elemental map of Figure 3I was similarly computed by picking pixels whose rate of change of signal going from camera lengths 44 mm to 55 mm > 25 and < 0, and going from camera lengths 55 mm to 69 mm was >12 and < 35. The rate of change of signal for endogenous cellular material from camera length 44 mm (Ɵ_1_ = 132 mrad) to camera length 55 mm (Ɵ_1_ = 106 mrad) is 17.9 ± 1.1; and from camera length 55 mm (Ɵ_1_ = 106 mrad) to camera length 69 mm (Ɵ_1_ = 84.1 mrad) is 8.0 ± 0.6. The endogenous cellular material map of Figure 3J was therefore calculated by picking pixels whose rate of change of signal going from camera lengths 44 mm to 55 mm < 25 and going from camera lengths 55 mm to 69 mm was <12. Figure 3K shows the Multicolor Z-contrast STEM image of DNA labeling by the Cu label (magenta) and the ER labeling by the Au label (cyan). Therefore, Multicolor EM using the Z-contrast STEM is a pseudo-chemical map utilizing differences in the signal (i.e. greyscale values in the image to create the color image), unlike the EELS/EFTEM technique which acquires true chemical maps by spectroscopic analysis.

**Figure 3.**
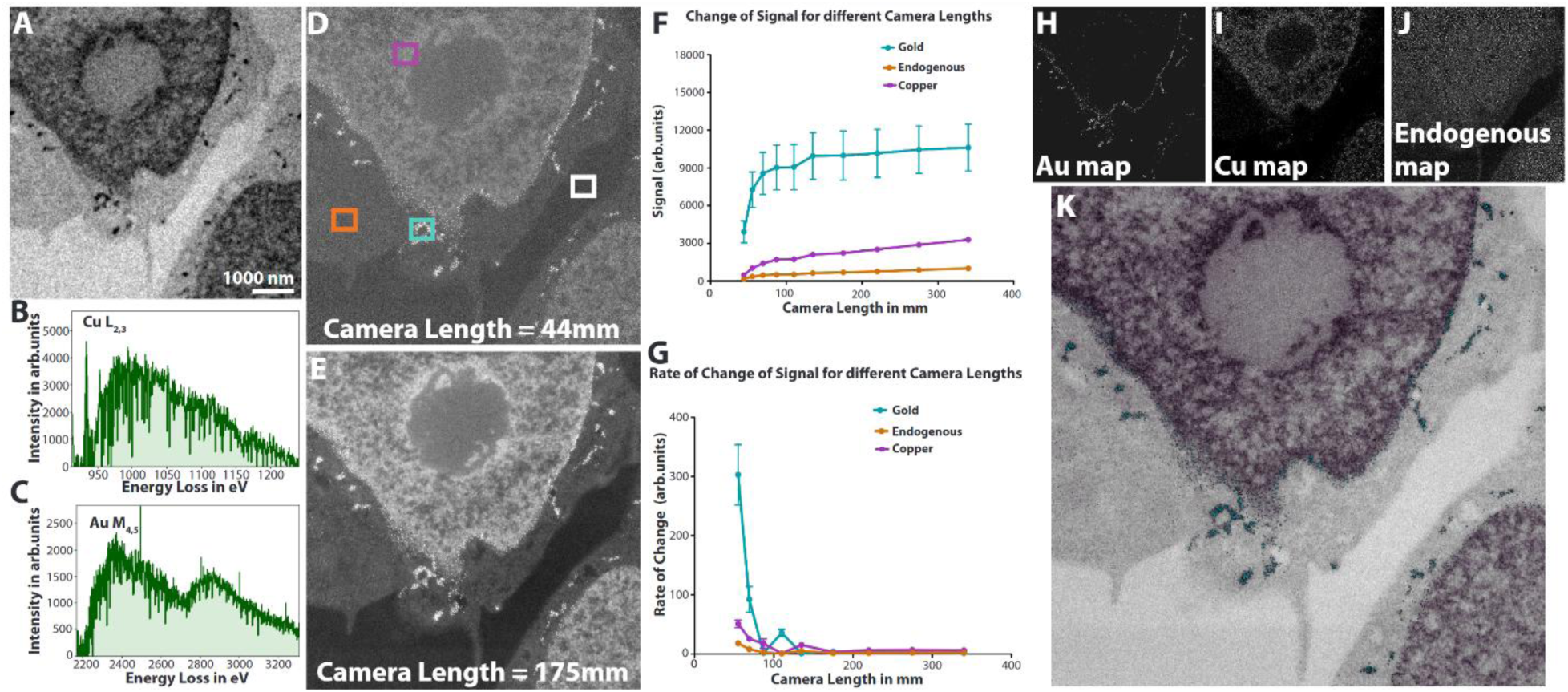
ColorSTEM datasets acquired on the HaaDF detector at different camera lengths (i.e. different scattering angles). The HEK cell samples were prepared by click-labeling of FeTAML-azide to EdU in DNA subsequently depositing Cu-DAB2, and ER expressing mSOG-KDEL depositing DAB enhanced with Au. No post-staining to avoid other metals. **A**, Conventional TEM image revealing Cu on the DNA and Au in the ER. **B**, EELS spectrum confirming the presence of Cu in the sample. **C**, EELS spectrum confirming the presence of Au in the sample. **D**, STEM HaaDF image collected at the smallest camera length of 44 mm i.e. the largest scattering semi-angle of 132 to 200 mrad. The blue, pink, and orange boxes indicate areas where signal from Au, Cu and endogenous cellular material were measured respectively, after normalizing for the signal from the resin. **E**, STEM HaaDF image collected at an intermediate camera length (175 mm) i.e. the scattering semi-angle of 33.2 to 200 mrad. The white colored box indicates a region that contains only resin; all the HaaDF STEM images were normalized to this signal at the respective camera length, by its subtraction. **F**, change in signal intensities for Au, Cu and endogenous cellular material as a function of camera length. **G**, rate of change in signal intensities, i.e. slope of the graph shown in **E1**, for Au and endogenous cellular material as a function of camera length. **H**, Au elemental map, calculated from the rate of change of Au signal with camera length. **I**, Cu elemental map, calculated from the rate of change of Cu signal with camera length. **J**, signal from endogenous material, calculated by the rate of change of endogenous signal with camera length. The STEM images collected at the three camera lengths, 44 mm, 55 mm and 69 mm would suffice to calculate the elemental maps shown in **H**, **I** and **J**. Although the graphs and trendlines have been calculated for larger camera lengths (44 mm to 340 mm) for sake of completeness of analysis, for real world colorSTEM maps a much smaller subset should be more than plentiful. **K**, Multicolor image with DNA labeled with Cu (magenta) and the ER labeled with Au (cyan).

To achieve MultiColorEM by Z-contrast in a single image and with high throughput we extended multiple labelled ‘Z-contrast’ samples to SEM Back Scatter Electron (BSE) imaging. BSE imaging in the SEM is analogous to STEM Z-contrast imaging. The BSE signal arises due to the incident electron having been elastically scattered multiple times by the atoms of the sample, the cumulative effect of these scattering events deviates the trajectory of the electron beam by > 90 degrees, resulting in electrons exiting the sample surface and captured by the BSE detector. The BSE signal shows a monotonic increase in signal with an increase in the atomic number (Z) of the target for electron beam energy > 5KV; and therefore, is also usually referred to as Z-Contrast or atomic number contrast ^41^. Figure 4 shows BSE contrast imaging in SEM of a Multicolor EM sample prepared by click-labeling DNA (with FeTAML-azide and EdU) in HEK cells, precipitating metal-free DAB conjugate that is subsequently labeled with Cu. Mitochondria were labeled by transfection with miniSOG-Mitomatrix, photosensitization of DAB and enhancement with Au. After fixation and no post staining, 100 nm thick sections on a regular TEM grid were imaged on a Zeiss stage biasing sample holder (Figure 4A), which can hold multiple sample stubs. The gold ribbon at the bottom, has electrical contacts to apply the desired uniform negative voltage on the sample. The samples used TEM grids in STEM-in-SEM kind of holders, and this was preferred over thin sections mounted on a silicon wafer to avoid contaminating signal from the underneath silicon substrate. A SEM sample stub was machined by drilling a hole (Figure 4B) to hold TEM grids, similar to STEM-in-SEM sample holders, to allow mounting on the Zeiss stage biasing sample holder.

**Figure 4.**
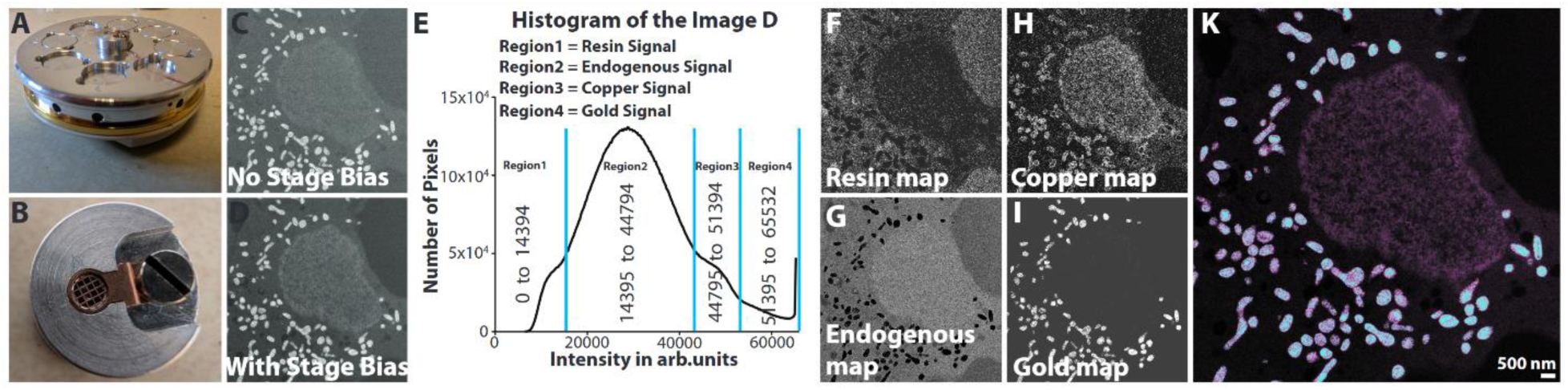
Multicolor Z-contrast SEM imaging. A, Zeiss stage bias sample holder, capable of holding multiple sample stubs. The gold ribbon at the bottom, has electrical contacts to apply the desired uniform negative voltage on the sample. B, SEM sample stub machined by drilling hole to hold TEM grids similar to STEM-in-SEM sample holders. C, BSD (Back-Scatter detector) image of Multicolor EM HEK cell sample prepared by click-reaction of FeTAML-azide to EdU in DNA, labeled with Cu-DAB2, and mitochondria expressing mSOG labeled with Au enhanced DAB, acquired at 5KV. D, BSD image of sample acquired at a beam energy of 7 KV and a deceleration of 2KV. E, histogram of the image in Figure 5B, showing 4 distinct regions. Segregating the image in 5B according to the marked 4 regions, the Z-Contrast signal from the sample is deconvolved into signal from: F, plastic resin. G, endogenous cellular material. H, specific Cu label on DNA and cross contamination on mitochondria. I, Au label on mitochondria. K, Multicolor SEM image with DNA labeled with Cu (magenta) and mitochondria labeled with Au (cyan).

The BSD image of the sample (Figure 4C) was acquired at 5 keV. Predictably, the Au signal from the mitochondria is very strong due to its high atomic number, and the Cu signal from the DNA is faint. When the electron beam interacts with the sample, it produces BSE electrons, whose energy varies from zero to E, where E is the incident energy of the beam. It also produces secondary electrons (SE), whose energy varies from 0 – 50 eV ^41^, but they are not energetic enough to produce a signal on the BSD. Unlike the BSE, the SE’s do not show any correlation with atomic number of the sample ^42^. However, if a negative stage bias is used, say −2 keV, then the SE will be accelerated towards the detector with an additional energy of 2 keV that is sufficient to create a signal on the BSD. Figure 4D shows such a BSD image of the above sample acquired at a beam energy of 7 keV and a deceleration of 2 keV. The Au signal from the mitochondria is still strong, due to both high BSE and SE signals but the signal from the Cu tag on the DNA is increased due to the high SE yield from Cu contributing to the signal. The histogram of the image in Figure 4D reveals four distinct regions and is marked in Figure 4E. By segregating the image in Figure 4D according to the signal in these four distinct regions, the Z-contrast signal from the sample can be deconvolved into (1) signal from the plastic resin (Figure 4F), (2) the endogenous cellular material (Figure 4G), (3) the Cu stain on the DNA, with some undesired cross contamination of the Cu stain on the mitochondria (Figure 4H), and (4) the Au signal on the mitochondria (Figure 4I). Figure 4K shows the Multicolor SEM image, with the colored (pseudo) chemical maps overlaid over the conventional SEM image.

Multicolor Z-contrast STEM imaging is a complimentary technique to EELS/EFTEM Multicolor EM, with the advantages including ease of use, fully automated acquisition, a relatively lower cost of required instrumentation, and minimal to no sample damage. Its applicability to SEM imaging is a further advantage that potentially could be applied to block-face imaging. The main disadvantage is that it is not a robust technique like EELS/EFTEM, which provides a true elemental/chemical map. However, both EELS/EFTEM and Multicolor Z-contrast STEM imaging are time consuming; EELS/EFTEM requires long exposures due to inherent low SNR. Multicolor Z-contrast STEM needs a beam shower routine at a lower magnification for 60 – 90 sec in between every acquisition to stabilize the sample from contamination buildup ^30, 34^. Both techniques are therefore not well suited for acquisition of large throughput datasets.

## ACKNOWLEDGEMENTS

We would like to thank David Mastronarde of University of Colorado, Boulder for his help with Serial EM scripting. This work was supported by NIH grant 5R01GM138780-04.

## Notes

### Competing Interest Statement

The authors have declared no competing interest.

## REFERENCES

1. Sanderson, M.J., Smith, I., Parker, P. & Bootman, M.D. Fluorescence Microscopy, Vol. Cold Spring Harb Protoc. 2014 Oct1;2014(10):pdb.top071795. (Springer, 2014).

2. Lichtman, J.W. & Conchello, J.A. Fluorescence microscopy. Nat Methods 2, 910–919 (2005).

3. Huang, B., Bates, M. & Zhuang, X. Super-resolution fluorescence microscopy. Annual review of biochemistry 78, 993–1016 (2009).

4. Adams, S.R. et al. Multicolor Electron Microscopy for Simultaneous Visualization of Multiple Molecular Species. Cell Chem Biol 23, 1417–1427 (2016).

5. Boassa, D. et al. Split-miniSOG for Spatially Detecting Intracellular Protein-Protein Interactions by Correlated Light and Electron Microscopy. Cell Chem Biol 26, 1407–1416 e1405 (2019).

6. Steinkellner, T. et al. Genetic Probe for Visualizing Glutamatergic Synapses and Vesicles by 3D Electron Microscopy. ACS Chem Neurosci 12, 626–639 (2021).

7. Sastri, M. et al. Sub-mitochondrial localization of the genetic-tagged mitochondrial intermembrane space-bridging components Mic19, Mic60 and Sam50. J Cell Sci 130, 3248-+ (2017).

8. Adams, S.R., et al. Fe-TAMLs as a new class of small molecule peroxidase probes for correlated light and electron microscopy. bioRxiv (2023).

9. Ramachandra, R. et al. Improving signal to noise in labeled biological specimens using energy-filtered TEM of sections with a drift correction strategy and a direct detection device. Microsc Microanal 20, 706–714 (2014).

10. Ramachandra, R. et al. Elemental mapping of labelled biological specimens at intermediate energy loss in an energy-filtered TEM acquired using a direct detection device. J Microsc 283, 127–144 (2021).

11. Scotuzzi, M. et al. Multi-color electron microscopy by element-guided identification of cells, organelles and molecules. Sci Rep-Uk 7 (2017).

12. Carter, C.B. & Williams, D.B. Transmission Electron Microscopy. (Springer, 2009).

13. Allen, L.J., D’Alfonso, A.J., Freitag, B. & Klenov, D.O. Chemical mapping at atomic resolution using energy-dispersive x-ray spectroscopy. Mrs Bull 37, 47–52 (2012).

14. Longo, P., Thomas, P., Aitouchen, A., Schaffer, B. & Twesten, R.D. Simultaneous EELS/EDS Composition Mapping at Atomic Resolution Using Fast STEM Spectrum-Imaging. Microscopy Today, 36–40 (2013).

15. Loane, R.F., Xu, P. & Silcox, J. Incoherent Imaging of Zone Axis Crystals with Adf Stem. Ultramicroscopy 40, 121–138 (1992).

16. Hartel, P., Rose, H. & Dinges, C. Conditions and reasons for incoherent imaging in STEM. Ultramicroscopy 63, 93–114 (1996).

17. Liu, J. & Cowley, J.M. High-Resolution Scanning-Transmission Electron-Microscopy. Ultramicroscopy 52, 335–346 (1993).

18. Pennycook, S.J. Z-Contrast Stem for Materials Science. Ultramicroscopy 30, 58–69 (1989).

19. Pennycook, S.J., Berger, S.D. & Culbertson, R.J. Elemental Mapping with Elastically Scattered Electrons. J Microsc-Oxford 144, 229–249 (1986).

20. Otten, M.T. High-Angle Annular Dark-Field Imaging on a Tem/Stem System. J Electron Micr Tech 17, 221–230 (1991).

21. Isaacson, M., Kopf, D., Ohtsuki, M. & Utlaut, M. Atomic Imaging Using the Dark-Field Annular Detector in the Stem. Ultramicroscopy 4, 101–104 (1979).

22. Howie, A. Image-Contrast and Localized Signal Selection Techniques. J Microsc-Oxford 117, 11–23 (1979).

23. Rutherford, E. The Scattering of alpha and beta Particles by Matter and the Structure of the Atom. Philos Mag 21, 669–688 (1911).

24. Crewe, A.V., Langmore, J.P. & Isaacson, M. Resolution and Contrast in the Scanning Transmission Electron Microscope. (1975).

25. Green, M.A., Sviland, L., Malcolm, A.J. & Pearson, A.D. Improved method for immunoperoxidase detection of membrane antigens in frozen sections. J Clin Pathol 42, 875–880 (1989).

26. Teclemariam-Mesbah, R., Wortel, J., Romijn, H.J. & Buijs, R.M. A simple silver-gold intensification procedure for double DAB labeling studies in electron microscopy. J Histochem Cytochem 45, 619–621 (1997).

27. Rae, J. et al. A robust method for particulate detection of a genetic tag for 3D electron microscopy. Elife 10 (2021).

28. Adams, S. et al. in Cell Chemical Biology, Vol. 23, Edn. Nov 17 1-11 (Cell Press, 2016).

29. Bushong, E.A., Martone, M.E., Jones, Y.Z. & Ellisman, M.H. Protoplasmic astrocytes in CA1 stratum radiatum occupy separate anatomical domains. J Neurosci 22, 183–192 (2002).

30. Egerton, R.F., Li, P. & Malac, M. Radiation damage in the TEM and SEM. Micron 35, 399–409 (2004).

31. Egerton, R.F. & Rauf, I. Dose-rate dependence of electron-induced mass loss from organic specimens. Ultramicroscopy 80, 247–254 (1999).

32. Tseng, Q.Z. et al. A new micropatterning method of soft substrates reveals that different tumorigenic signals can promote or reduce cell contraction levels. Lab Chip 11, 2231–2240 (2011).

33. Mastronarde, D.N. Automated electron microscope tomography using robust prediction of specimen movements. J Struct Biol 152, 36–51 (2005).

34. Mitchell, D.R. Contamination mitigation strategies for scanning transmission electron microscopy. Micron 73, 36–46 (2015).

35. Sousa, A.A., Hohmann-Marriott, M.F., Zhang, G. & Leapman, R.D. Monte Carlo electron-trajectory simulations in bright-field and dark-field STEM: Implications for tomography of thick biological sections. Ultramicroscopy 109, 213–221 (2009).

36. Jones, L. Quantitative ADF STEM: acquisition, analysis and interpretation. Iop Conf Ser-Mat Sci 109 (2016).

37. Ishikawa, R., Lupini, A.R., Findlay, S.D. & Pennycook, S.J. Quantitative Annular Dark Field Electron Microscopy Using Single Electron Signals. Microsc Microanal 20, 99–110 (2014).

38. Krivanek, O.L. et al. Atom-by-atom structural and chemical analysis by annular dark-field electron microscopy. Nature 464, 571–574 (2010).

39. Treacy, M.M.J. & Gibson, J.M. Coherence and Multiple-Scattering in Z-Contrast Images. Ultramicroscopy 52, 31–53 (1993).

40. Langlois, C., Wang, Z., Pearmain, D., Ricolleau, C. & Li, Z. in Journal of Physics: Conference Series, Vol. 241 012043 (2010).

41. Goldstein, J.I. et al. Scanning Electron Microscopy and Microanalysis. (Springer, 2003).

42. Lin, Y. & Joy, D.C. A new examination of secondary electron yield data. Surface and Interface Analysis 37, 895–900 (2005).

